# *Sinorhizobium meliloti* FcrX coordinates cell cycle and division during free-living growth and symbiosis

**DOI:** 10.1101/2023.03.13.532326

**Authors:** Sara Dendene, Shuanghong Xue, Quentin Nicoud, Odile Valette, Angela Frascella, Anna Bonnardel, Romain Le Bars, Mickaël Bourge, Peter Mergaert, Matteo Brilli, Benoît Alunni, Emanuele G. Biondi

**Author notes:** co-last authors (Benoît Alunni, Emanuele G. Biondi).

## Abstract

*Sinorhizobium meliloti* is a soil bacterium that establishes a symbiosis within root nodules of legumes (*Medicago sativa*, for example) where it fixes atmospheric nitrogen into ammonia and obtains in return carbon sources and other nutrients. In this symbiosis, *S. meliloti* undergoes a drastic cellular change leading to a terminal differentiated form (called bacteroid) characterized by genome endoreduplication, increase of cell size and high membrane permeability. The bacterial cell cycle (mis)regulation is at the heart of this differentiation process. In free-living cells, the master regulator CtrA ensures the progression of cell cycle by activating cell division (controlled by the tubulin-like protein FtsZ) and simultaneously inhibiting supernumerary DNA replication, while on the other hand the downregulation of CtrA and FtsZ is essential for bacteroid differentiation during symbiosis, preventing endosymbiont division and permitting genome endoreduplication. Little is known in *S. meliloti* about regulators of CtrA and FtsZ, as well as the processes that control bacteroid development. Here, we combine cell biology, biochemistry and bacterial genetics approaches to understand the function(s) of FcrX, a new factor that controls both CtrA and FtsZ, in free-living growth and in symbiosis. Depletion of the essential gene *fcrX* led to abnormally high levels of FtsZ and CtrA and minicell formation. Using multiple complementary techniques, we showed that FcrX is able to interact physically with FtsZ and CtrA. Moreover, its transcription is controlled by CtrA itself and displays an oscillatory pattern in the cell cycle. We further showed that, despite a weak homology with FliJ-like proteins, only FcrX proteins from closely-related species are able to complement *S. meliloti fcrX* function. Finally, deregulation of FcrX showed abnormal symbiotic behaviors in plants suggesting a putative role of this factor during bacteroid differentiation. In conclusion, FcrX is the first known cell cycle regulator that acts directly on both, CtrA and FtsZ, thereby controlling cell cycle, division and symbiotic differentiation.

## INTRODUCTION

*Sinorhizobium meliloti* belongs to the class of *Alphaproteobacteria* and is known for its dual lifestyle: as a free-living bacterium in the soil and as a symbiotic endophyte within legumes of the genera *Medicago*, *Melilotus* and *Trigonella* ^1^. Free-living *S. meliloti* cells thrive in the soil and divide asymmetrically to produce two physiologically and morphologically different cell types (Figure 1A): a smaller cell, unable to replicate its DNA and a larger cell able to replicate its DNA once per cell cycle. The small cell first has to differentiate into the large cell type, upon suitable nutrient conditions, before it can undergo genome replication ^2^.

**Figure 1:**
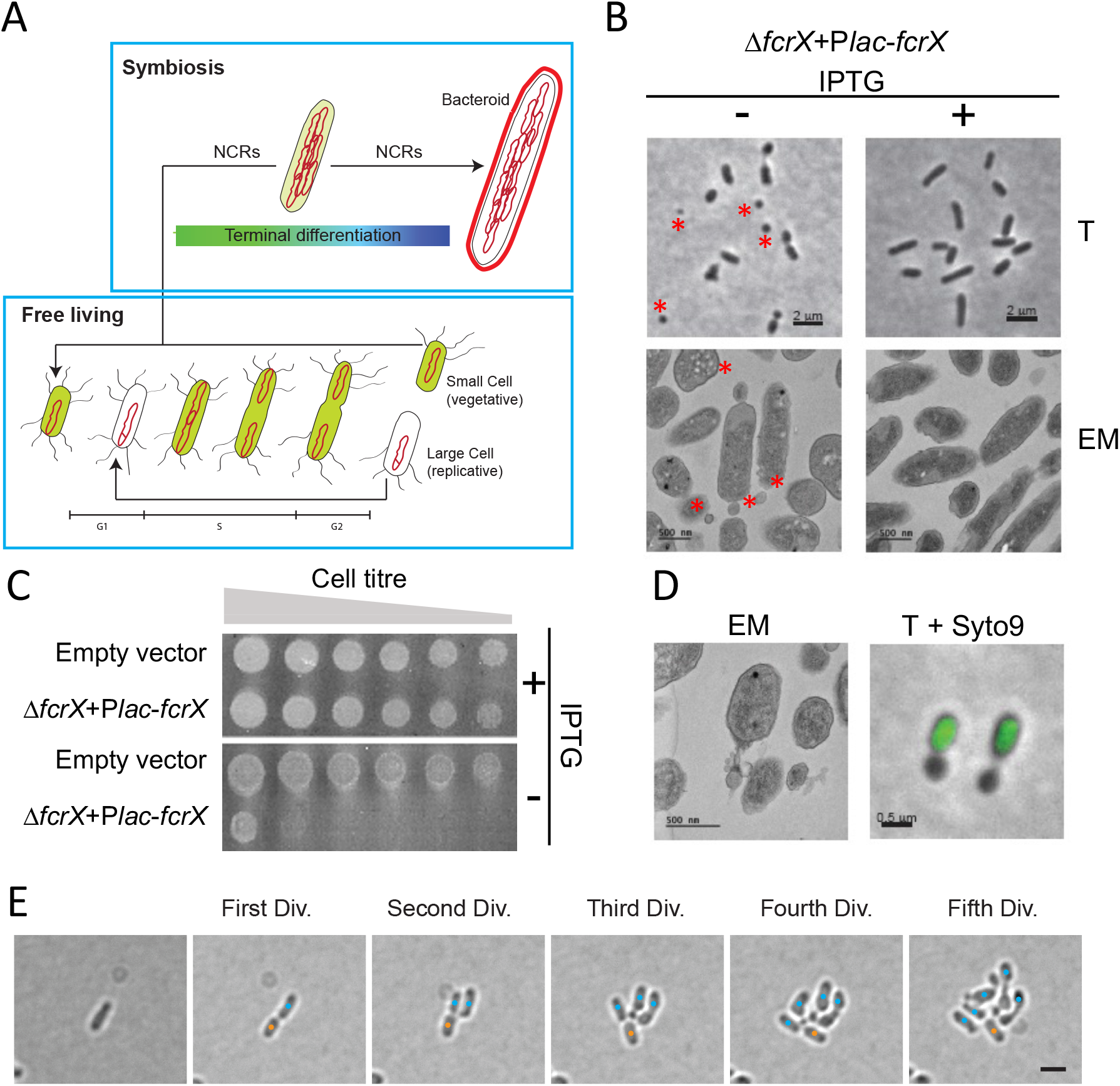
*fcrX* is an essential gene in *S. meliloti*. (A) Scheme representing the cell cycle progression of *S. meliloti* in free-living condition and in symbiosis with legume plants. The green color refers to CtrA concentration, which is low at the beginning of S-phase and after cell division in large cells. During bacteroid differentiation CtrA is removed in order to induce morphological changes. (B) Transmission contrast (T) and electron microscopy (EM) of a depletion strain of *fcrX* in presence and absence of the inducer (IPTG). In particular, in no IPTG conditions, the depletion of *fcrX* is causing the formation of small cells (red asterisks). (C) Viability test on *S. meliloti* containing empty plasmid and a depletion strain of *fcrX*. Cells of *fcrX* depletion strain or wild type strain carrying an empty plasmid were grown with IPTG and then washed before plating. From left to right, non-diluted to 1/10^6^ diluted cell suspensions were spotted on an agar plate with or without IPTG. (D) Electron microscopy of a depletion strain of *fcrX* (EM) and overlay of transmission contrast (T) and epifluorescence microscopy of a depletion strain of *fcrX* labeled with Syto9. (E) Time lapse microscopy on a depletion strain of *fcrX*. Orange dots represent the mother cells and the blue dots represent the daughter cells. Only daughter cells are able to produce abnormal mini cells. Scale bar corresponds to 2 μm.

To ensure a normal progression of the cell cycle, it has to be tightly regulated. In *Alphaproteobacteria* the response regulator CtrA plays a central role in this process by binding to the DNA, mostly activating or inhibiting transcription of more than a hundred target genes ^3,4^. The function of CtrA has been mainly studied in *Caulobacter crescentus*, an aquatic bacterium that divides asymmetrically, as *S. meliloti*, giving two morphologically and physiologically different daughter cells ^5^; a stalked cell, able to replicate immediately after division, and a non-replicative motile cell that is blocked at the G1 phase. CtrA is under a strict control, as its levels oscillate during the cell cycle, reaching a maximum in two moments: i. at the G1 phase (motile cell), when it inhibits DNA replication by binding to specific sites that prevent origin binding of DnaA, the initiation factor of DNA replication, and ii. at the end of S-phase/G2, when CtrA also activates the cell division process ^6^. To permit this pattern, regulation mechanisms at the transcriptional and post translational (by phosphorylation and proteolysis) levels are involved ^7,8^. An homolog of CtrA is present in *S. meliloti*, where it has a similar function ^9,10^. In *S. meliloti*, CtrA inhibits indirectly the DNA replication by a yet-unknown process and activates the cell division by repressing the transcription of the *minCDE* system ^10^, which ultimately inhibits the polymerization of the tubulin-like FtsZ responsible of cell constriction at the septum ^11^. The FtsZ protein is composed by a N terminal core region, containing a GTPase domain, involved in the polymerization activity, and a C terminus responsible for the interaction to other actors^12^. In *S. meliloti*, FtsZ is present in two copies, FtsZ1 and FtsZ2, however only FtsZ1 is essential for the Z ring formation, while the second copy lacks the C-terminal domain and its deletion is not lethal ^13^. As many cell cycle actors, FtsZ1 is expressed at the predivisional phase of the cell cycle ^14^.

In symbiosis with *Medicago* plants, *S. meliloti* colonizes special root organs, called nodules. There, it fixes atmospheric nitrogen into ammonium that is assimilated by the plant, while it receives in return dicarboxylic acids and other nutrients ^15^. In nodules, after a stage of multiplication, *S. meliloti* undergoes a drastic cellular change into a terminally differentiated form called bacteroid. This process takes place intracellularly, inside the nodule plant cells, where differentiated bacteroids are characterized by genome endoreduplication, cell enlargement and high membrane permeability ^16^. Previous studies have shown an implication of the bacterial cell cycle regulation in this differentiation process (Pini et al., 2013-2015; Kobayashi et al., 2009). Indeed, CtrA and FtsZ are absent in bacteroids ^17,18^ and mutants that overexpress CtrA are characterized by a symbiotic defect ^17^. Interestingly, the depletion of *ctrA* in *S. meliloti*, similarly to *C. crescentus*, leads to a cell elongation/enlargement and endoreduplication phenotype that is strikingly similar to bacteroids formed in nodules. The mechanism leading to the downregulation of CtrA and FtsZ in bacteroids is not known yet but studies showed the involvement of plant-produced Nodule-specific Cysteine-Rich (NCR) peptides ^19,20^ (Figure 1A). Legumes such as *Medicago truncatula* produce a wide spectrum of NCR peptides (about 600) that are implicated in the disruption of several cellular processes including the cell cycle, thus resulting in the terminal differentiation of *S. meliloti* (Mergaert *et al.*, 2006; Van de Velde *et al.*, 2010; Alunni & Gourion, 2016). Indeed, the treatment of a wild type strain of *S. meliloti* with the NCR247 showed a down regulation of CtrA and its regulon. Farkas and colleagues also highlighted a physical interaction between this NCR and FtsZ (Farkas *et al*., 2014). Overall these data strongly suggest that the regulation of bacterial cell cycle and cell division are playing a major role in the symbiosis process ^21^.

Here we characterized the role of a new cell cycle regulator, named FcrX, elucidating its role with respect to CtrA and FtsZ1 and FtsZ2, its regulation by transcription, its conservation across the class *Alphaproteobacteria* and finally, its role during symbiosis.

## RESULTS

### The *fcrX* gene is essential and controls cell cycle in *Sinorhizobium meliloti*

The *fcrX* gene (SMc00655, 351 bp ORF coding for a 116 aa predicted protein) is adjacent to *ctrA* in the *S. meliloti* chromosome in a head-to-head orientation, with a *ca*. 500 bp long intergenic sequence that controls the transcription of both genes. The *fcrX* gene appears essential in free-living conditions, based on Tn-seq results (Figure S1A). In order to study the function of *fcrX*, we constructed a depletion strain by deleting the chromosomal copy of *fcrX* and expressing an extra copy of *fcrX* on a plasmid under the control of an IPTG-inducible *lac* promoter ^17,22^. This conditional depletion strain of FcrX grew best in a medium supplemented with 100 μM IPTG (Figure S1B) and showed upon removal of IPTG dramatic growth defects (Figure 1B and Figure S1B), as well as a reduced capacity to form colonies (Figure 1C), indicating that *fcrX* is essential for the viability of *S. meliloti*. To gain more insights in FcrX function(s), we investigated the FcrX-depleted cells by microscopy and flow cytometry. Interestingly, the absence of FcrX induced the production of DNA-free small cells, referred to as minicells, as revealed by DNA staining by Syto9 and microscopy observation (Figure 1D). The accumulation of minicells in an FcrX-depleted suspension was confirmed by DNA/membrane double staining with DAPI/Potomac Gold dyes and flow cytometry analysis (Figure S2-S3B, 14.99% of total events with IPTG against 36.91% of total events without IPTG). Thus upon depletion of *fcrX*, an accumulation of small cells with no DNA was observed. We monitored the depletion of FcrX by time-lapse microscopy in order to understand the development of minicells at the cellular level. Minicells tend to form only in daughter cells (small cell) while the mother (large) cell remains able to divide efficiently producing daughter cells, until two additional cycles and then completely stops dividing (Figure 1E). This minicell phenotype can be interpreted as the consequence of an imbalanced cell cycle, with an excess of division and a block of DNA replication, suggesting that FcrX may coordinate these two processes in *S. meliloti*. Consistently, the absence of FcrX led to increased levels of CtrA, FtsZ1 and FtsZ2 proteins, as shown by western blot analysis (Figure 2A), indicating that FcrX indeed negatively controls the accumulation of CtrA and FtsZ1/2. This result strengthens our hypothesis of the implication of FcrX in the regulation of cell cycle and cell division.

**Figure 2:**
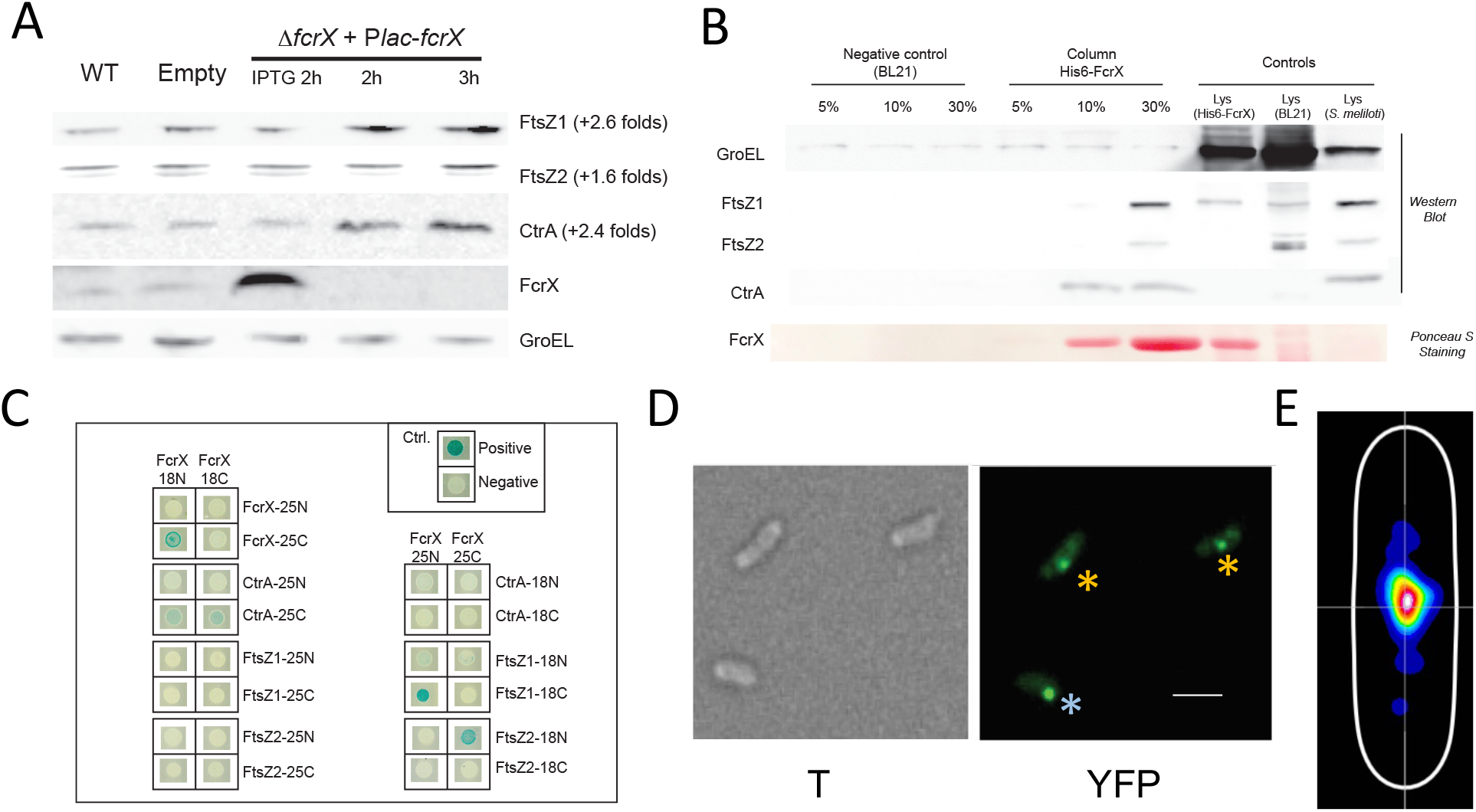
FcrX down regulates and interacts directly with the master regulator CtrA and the Z ring proteins (FtsZ1, FtsZ2). (A) Western blot using Anti-FtsZ, CtrA, FcrX and GroEL on the *fcrX* depletion strain in comparison with wild type (WT) and a strain containing the empty vector used in the *fcrX* depletion strain (Empty). For the depletion strain of *fcrX*, a depleted culture was reincubated with IPTG for 2h (IPTG 2h) or without for 2h (2h) or 3h (3h). (B) Affinity column western blot using CtrA, FcrX and FtsZ antibodies. Percentages represent imidazole concentration (see Materials and Methods for details). Membrane stained with Ponceau Red shows His6-FcrX. (C) Bacterial Two hybrid (BACTH) testing FcrX interaction with FtsZ1, FtsZ2, CtrA and itself. Upper right box shows negative and positive controls provided by the supplier. 25 and 18 are the two subunits of adenylate cyclase of the BACTH (see Materials and Methods). (D) Functional FcrX-YFP C-terminal fusion observed by transmission (T) and epifluorescence microscopy (EM) in a *fcrX* deletion genetic background (see text for details). Bar corresponds to 1 μm. Cells showing mid-cell localization are marked with an orange asterisks, while the cell with polar localization is marked with a blue asterisk. (E) Heatmap of YFP-FcrX subcellular localization in a synchronized cell population (predivisional phase, 150 minutes). Analysis performed on >300 cells (56 with foci).

### FcrX interacts with CtrA and FtsZ1/2

The 3D structure prediction of FcrX performed with the Alphafold 2 algorithm showed that the FcrX protein may be composed of two alpha helices in a coiled coil configuration (Figure S4). The structure did not reveal the presence of a DNA binding domain, suggesting that its putative mode of action on CtrA and FtsZ1/2 may be by a direct interaction at the protein level. To verify this hypothesis, an affinity column experiment was realized using His6-FcrX loaded nickel columns in order to identify potential FcrX-interacting proteins. The presence of FtsZ1/2 and FcrX itself, together with other proteins listed in Table S1 was detected in the affinity column eluate by mass spectrometry (Figure S5). The presence of CtrA on the other hand was only detected by western blot (Figure 2B), as its low abundance makes it hardly detectable by mass spectrometry. In order to confirm this result, we carried out a bacterial two-hybrid (BACTH) experiment using the *Escherichia coli* carrier (Figure 2C) (Karimova *et al.*, 1998). FcrX, CtrA and FtsZ1/2 proteins were fused to domains 18 and 25 of the adenylate cyclase from *Bordetella pertussis* in C- and N-terminal orientations and all possible combinations of prey and bait were introduced in the *E. coli* strain HB101. In this system, FcrX interacts with CtrA and with both copies of FtsZ (1 and 2). FtsZ2 lacks the C-terminal domain that is usually implicated in protein-protein interactions and recruitment of FtsZ partners ^23^. Thus, the observed interaction between FcrX and FtsZ2 may be explained by either an unusual interaction between the N-terminal domain or by the presence of a functional full-length FtsZ homolog in *E. coli* (FtsZ_Ec_), which could promote the formation of a FcrX-FtsZ_Ec_-FtsZ2 ternary complex. We also tested the interaction of FcrX with itself to verify the possible dimerization of the protein, as observed with the affinity column. As shown in the (Figure 2C) a strong FcrX-FcrX interaction was observed, which is consistent with a putative oligomeric FcrX structure.

### Subcellular localization of FcrX and its interactors

To gain more insights about the function of FcrX we decided to investigate its subcellular localization and whether it colocalizes with its interactors, FtsZ being shown to have a mid-cell localization and CtrA potentially being polarly localized in *S. meliloti* as it has been shown previously in *C. crescentus* ^24,25^. We constructed a strain expressing a C-terminal translational fusion of FcrX with a yellow fluorescent protein (YFP) and further deleted the chromosomal copy of *fcrX* by transduction using a Δ*fcrX*::tetR M12 phage lysate. This approach was used in order to avoid all possible competitions of the wild-type FcrX with the tagged version and it confirmed that the fusion protein retained its function. FcrX localized at the septum, as observed by epifluorescence microscopy (with a less frequent localization at the cell pole, presumably after cell division) (Figure 2D). Image analysis in predivisional phase cells confirmed that FcrX is localized at the septum in the majority of these cells (Figure 2E). We wanted to correlate this result with the subcellular localization of the interactors over the cell cycle. Therefore, we constructed a C-terminal translational fusion of FtsZ1 with a cyan fluorescent protein (CFP), which despite the loss of its functionality retained its native localization ^24^. Using a similar approach, we tagged FtsZ2 with CFP. CtrA localization investigation appeared more complex as N- or C-terminal tags strongly stabilized its levels, which is a lethal condition in *S. meliloti* ^10^. Therefore, we added a sequence coding for the last 15 amino acids of CtrA that constitute the degradation motif, downstream of CFP to ensure the timely degradation of the fusion protein by proteolysis, then deleted the chromosomal copy of *ctrA* by transduction using a *DctrA*::tetR M12 phage lysate. Tagged proteins were observed under the microscope. FtsZ1 and FtsZ2 showed as expected a midcell localization while CtrA showed a localization at the cell pole, consistently with FcrX localization (Figure S6). Together, these results indicate that FcrX may interact with both CtrA and FtsZ1/2 *in vivo* but at distinct subcellular localizations.

### Transcription of *fcrX* is positively regulated by CtrA

In order to define and characterize the *fcrX* promoter, we made a promoter deletion analysis by introducing in *S. meliloti* wild-type cells an extra copy of *fcrX* with different putative promoter lengths located on a plasmid downstream of the IPTG-regulated *lacZ* promoter (Figure S7). Then, the obtained clones were transduced using a phage M12 lysate produced from the strain *DfcrX*::tetR + Plac-*fcrX* and selected in the presence or absence of IPTG. All clones should give viable colonies with IPTG while only the constructs containing a functional promoter of *fcrX* should support viability without IPTG (Figure S7). Constructs with *PfcrX-fcrX* fragments containing at least the region between −1/−287 were viable, while a shorter promoter fragment (−1/−224) was not active, suggesting that the promoter region of *fcrX* resides within the −287 region and that the −224/−287 contains a critical promoter element.

To confirm these results, we used the plasmid pOT1em ^26^, which contains genes encoding mCherry and EGFP in opposite directions. Several derivatives of the *fcrX* promoter, described in Figure S7, as well as the full intergenic region between *fcrX* and *ctrA*, were cloned between the ATGs of mCherry and EGFP. The full intergenic region between *fcrX* and *ctrA* was able to express both mCherry (*fcrX*) and EGFP (*ctrA*) (Figure 3A). Analysis of the other constructs containing different versions of the upstream region of *fcrX* confirmed that only clones with at least −1/−287 expressed mCherry (*fcrX*) (data not shown). The *ctrA* gene is also controlled by the same intergenic region, suggesting that these two genes may share the same transcriptional regulation. The analysis of the region −224/−287 revealed several interesting characteristics. First, in this region the *ctrA*P1 promoter ^27^ and the estimated *pfcrX* are overlapping with RNA-seq determined putative TSSs (transcriptional start site) separated only by 20/25 bp (Figure S7). Second, the promoter of *fcrX* contains a CtrA binding box within the 50 bp region upstream the putative TSS ^4,10^. These observations suggest that the activation of the *fcrX* promoter may depend on CtrA. In order to test this hypothesis, we mutated the CtrA-binding box by replacing the 5’-TTAA-3’ half box with 5’-GCGC-3’ in a pOT1em plasmid carrying the full intergenic region between *ctrA* and *fcrX* (Figure 3A). It was shown that this mutation prevents the fixation of CtrA on the box and therefore affects its transcriptional activity on the downstream gene (Figure 3A) (ref). Microscopy observation showed that the strain containing the mutated CtrA-binding box didn’t express the mCherry fluorescence, implicating that CtrA may activate *fcrX* transcription by binding to its box. On the other hand, the mutated CtrA binding box didn’t change the EGFP expression noticeably, excluding the implication of this box in the regulation of *ctrA* transcription (Figure 3A).

**Figure 3:**
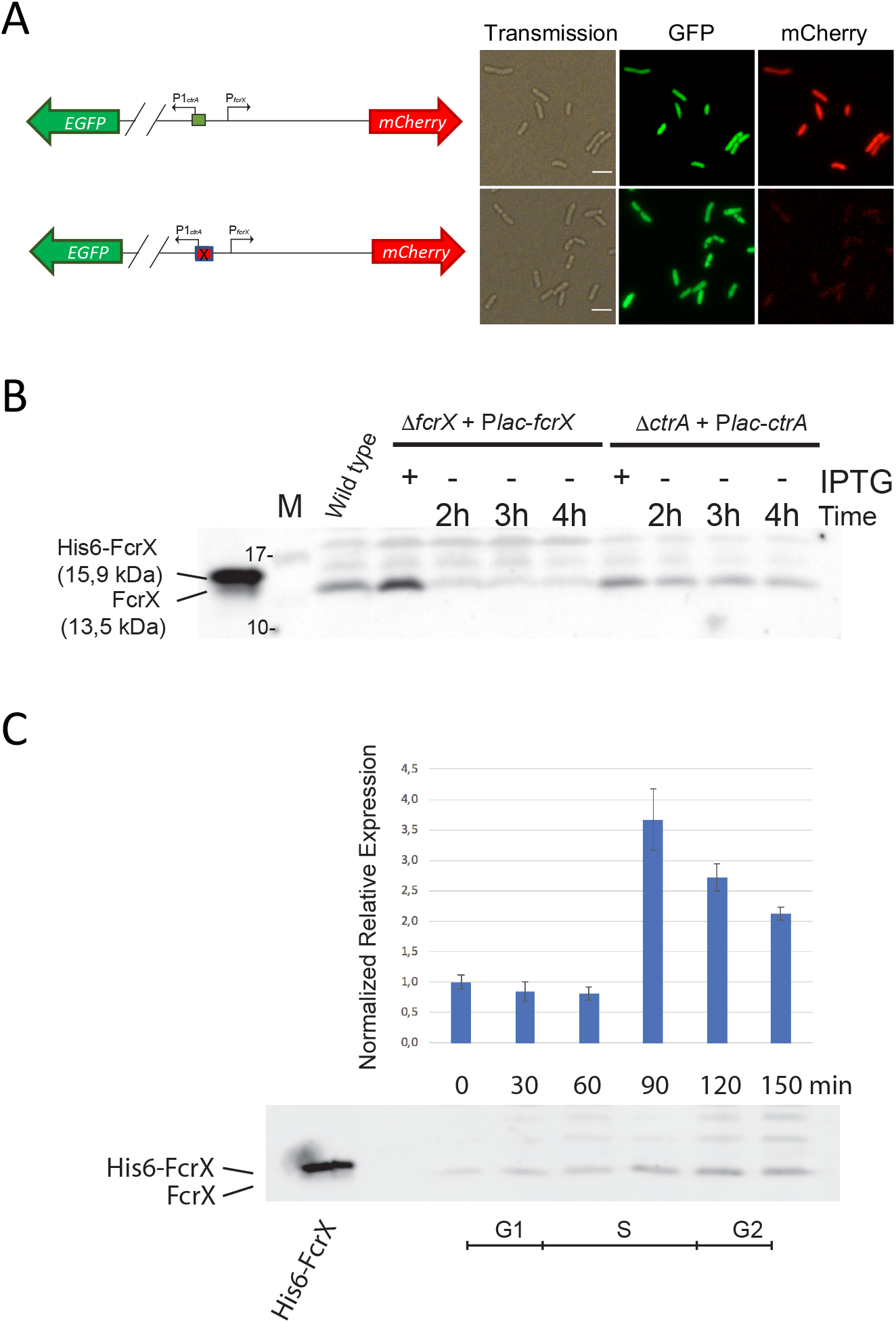
FcrX is cell cycle regulated and its transcription depends on CtrA. (A) *S. meliloti* strain containing the intergenic region between *fcrX* and *ctrA* fused with mCherry and EGFP, respectively. Lower panels correspond to the same intergenic region mutated in the CtrA box (see text for details). This mutation doesn’t affect the expression of *ctrA* but it completely abolishes the expression of *fcrX*. (B) Western blot using Anti FcrX antibodies using FcrX depletion and CtrA depletion samples. First lane is purified His6-FcrX. M = Marker (sizes are reported). (C) qRT-PCR of *fcrX* and Western blot using anti-FcrX antibodies on a synchronized population of *S. meliloti*. Bottom part represents a timeline of cell cycle phases.

To confirm the positive regulation of *fcrX* expression by CtrA, we tested the steady state levels of FcrX upon depletion of CtrA by western blot using antibodies directed against FcrX. In the absence of CtrA, we observed a significant decrease of FcrX (Figure 3B). Since CtrA is a DNA-binding protein implicated in transcriptional regulation, we also performed a qRT-PCR in the same conditions. Consistently, *fcrX* expression decreased upon CtrA depletion, confirming the previous observation (Figure S8). These results build up a regulatory model involving a negative feedback loop between CtrA and FcrX.

Cell cycle regulators are known to be dynamically regulated over the cell cycle, consistent with the oscillatory nature of this biological phenomenon. Because FcrX is closely linked to cell cycle regulation, we wanted to check whether FcrX was subject to oscillation. Therefore, a synchronization of a wild type culture of *S meliloti* was made as described before (Figure 3C) ^14^. Samples were recovered every 30 minutes over a full cell cycle and the corresponding total RNA was used to perform a qRT-PCR analysis while cell lysates were used for Western blot experiments using antibodies directed against FcrX. Both transcription and translation of FcrX increased at 90 min of the cell cycle, meaning that FcrX is not only a regulator of the cell cycle but it is also subject to cell cycle oscillation. These data are consistent with previous analyses on the ensemble of cell cycle regulated genes ^14^.

### FcrX is essential for the establishment of the legume symbiosis

The terminal bacteroid differentiation that *S. meliloti* undergoes during its interaction with *Medicago* plants involves a remodeling of the cell cycle and its regulatory network, prompting us to test the involvement of FcrX in the symbiotic process. We tested the importance of FcrX by inoculating *M. sativa* plants with the FcrX depletion strain and watered the plants with a range of concentrations of IPTG in order to obtain different levels of FcrX expression. We also inoculated plants with the wild type strain of *S. meliloti* carrying an empty plasmid as a reference. At 28 days post inoculation (dpi) we checked nodule colonization using confocal microscopy (Figure 4A), measured the plant dry weight (Figure 4B) and assessed bacteroid differentiation by measuring the bacterial DNA content with flow cytometry (Figure S9). Plant weight decreased when FcrX expression was reduced with respect to the condition with the highest IPTG concentration (Figure 4B). The confocal microscopy images showed a clear nodule colonization defect, with much less plant cells containing bacteroids in the plants watered with 0μM, 10μM and 100μM IPTG compared to the plants watered with 1mM IPTG and the references, implying that FcrX is essential to the establishment of a fully functional symbiosis. However, the nodules from the condition watered with 1mM IPTG were still less colonized and the bacteroids did not show a similar extent of terminal differentiation as observed in the reference conditions, an observation that can be explained by the IPTG-regulated expression of FcrX which may not mimic the natural cell cycle-regulated expression. Considering our results showing that FcrX inhibits the accumulation of CtrA, FtsZ1 and FtsZ2 (Figure 2A), we wondered if an overexpression of FcrX may promote CtrA and FtsZ1/2 disappearance inside nodule cells and thereby boost the symbiotic process. To do so, we inoculated plants of *M. sativa* with a *S. meliloti* strain containing a second copy of *fcrX* expressed on a pSRK plasmid under the control of the *Plac* promoter, which has a weak transcriptional leakage in *S. meliloti*, even in the absence of IPTG ^22^. The upregulation of FcrX resulted in a significant plant biomass gain at 28 dpi, as compared to the controls (Figure 4C). However, no increase in the bacteroid differentiation level was noticed, suggesting that the increase in symbiotic efficiency is likely related to the speed up of the differentiation process rather than to a larger extent of differentiation (data not shown). Finally, we investigated whether fully differentiated bacteroids contained FcrX by using anti-FcrX antibodies. Unlike CtrA and FtsZ1/2 that are absent from mature bacteroids, FcrX remained detectable (Figure 4D), supporting a role of FcrX during the establishment of bacteroid differentiation and maintenance.

**Figure 4:**
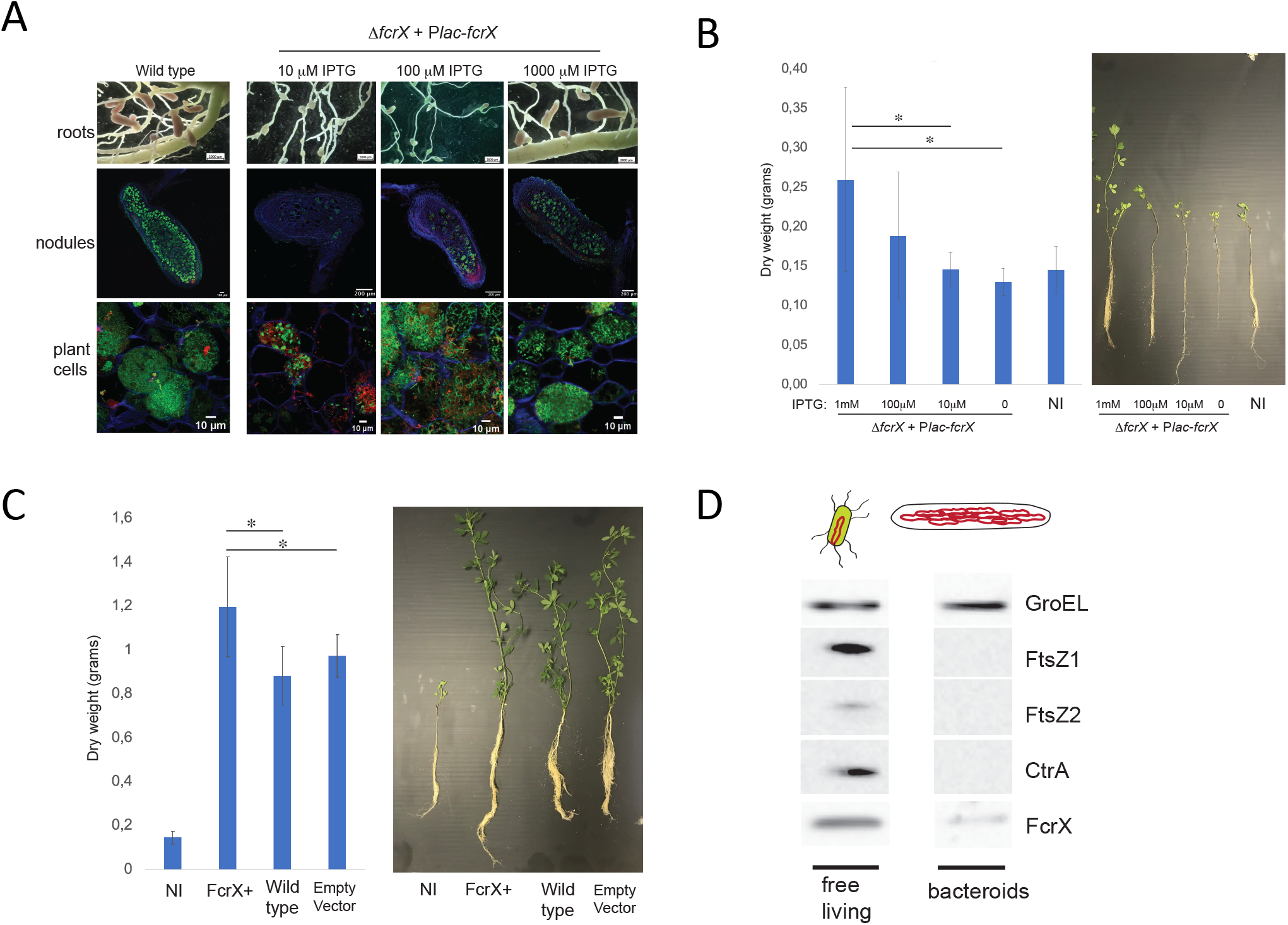
FcrX is important for symbiosis. (A) Nodules of 42 dpi plants infected by wild type (left row) and the depletion strain of *fcrX*, in different IPTG-watering conditions, were photographed (upper panels) and then sectioned and stained with Calcofluor, Syto9 and IP as explained in materials and methods (lower panels at high magnification levels). (B) Dry biomass per plant (left panel) and aspect (right panel) of 42 dpi *M. sativa* infected by wild type and the depletion strain of *fcrX* at different IPTG concentrations. Asterisks correspond to significant differences (More than 20 plants for each condition, less than P<0,05, Kruskall Wallis test). NI = Non inoculated. (C) Dry biomass per plant (left panel) and aspect (right panel) of 42 dpi *M. sativa* infected by wild type strain containing an empty vector (empty), wild type and a strain expressing an extra copy of *fcrX* (FcrX+). Asterisks correspond to significant differences (More than 20 plants for each condition, less than P<0,05, Kruskall Wallis test). NI = Non inoculated. (D) Western blot using FcrX, FtsZ, CtrA and GroEL antibodies on free living and bacteroid cells.

### FcrX is a conserved factor in several species of the class *Alphaproteobacteria*

We further wanted to investigate the presence and functionality of FcrX in other bacteria. First, the *S. meliloti* FcrX protein sequence was used to search in the Microbes Online database ^28^ for similar proteins. Orthologs defined by Bidirectional Blast Hit (BBH) of FcrX were found in several rhizobia (*Bradyrhizobium japonicum*, *Rhizobium leguminosarum*), the human pathogen *Brucella abortus* and the phytopathogen *Agrobacterium vitis*. Except for *A. vitis* for which no information is available, previous Tn-seq data on all those species revealed that the putative FcrXs were all essential in the organism of origin ^29–31^. Those orthologs were tested for their capacity to complement *fcrX* deletion in *S. meliloti*. We also tested a distant homolog of *fcrX* from *C. crescentus*, which codes for the flagellar protein FliJ, located in the same genomic context as *fcrX*. Interestingly, the *fcrX*-*ctrA* synteny is widely preserved among these bacteria suggesting a possible conservation of *fcrX* function (Figure 5A). In order to test our hypothesis, we constructed *S. meliloti* strains expressing a copy of these orthologs and we deleted by transduction the chromosomal copy of *fcrX*. Results of this experiment showed that the expression of the orthologs from *R. leguminosarum* and *B. abortus* were able to support *S. meliloti* growth in a *fcrX* deletion background, implying functional conservation of FcrX. A western blotting experiment was carried out confirming that the overexpression of these orthologs is able to down regulate the accumulation of FtsZ and CtrA (Figure S10). However, the orthologs from *B. japonicum*, *A. vitis* and the FliJ homolog from *C. crescentus* were not able to complement the *fcrX* deletion. This functional complementation analysis, although still limited in the number of tested species, revealed that from a functional point of view each FcrX has acquired specific functions independently from the phylogenetic distance from *S. meliloti*, as for example *A. vitis* (a species close to *S. meliloti*) is not able to complement while the more distant *B. abortus* does. In order to have a broader view of FcrX conservation, we searched for FcrX/FliJ orthologs/homologs in other alphaproteobacterial species (Figure 5B, Table S2). Significantly similar sequences to either the *S. meliloti* FcrX or the *C. crescentus* FliJ, were only found in the *Caulobacterales* and *Rhizobiales*. However, the two queries retrieved sequences in complementary sets of species; for example, in *C. crescentus* FcrX retrieved no results (Table S2). This can be consequent to a functional diversification of the same ancestral gene in the *Caulobacterales* and *Rhizobiales*, which is however difficult to demonstrate because of the short length and variability of these sequences, even if a phylogenetic tree of all homologs found seems to suggest an orthology relationship as the topology is largely congruent with the RecG tree (data not shown). In parallel we checked the distance between *fcrX*/*fliJ* and *ctrA* genes, discovering that the *ctrA* gene is in proximity to *fcrX* orthologs (or *fliJ* in *Caulobacterales*), and are often transcribed from the same intergenic region. Finally, we observed that orthologs able to complement the *S. meliloti fcrX* deletion belong to the phylogenetic group of *Brucella*-rhizobia (excluding bradyrhizobia). However, the functional complementation of *fcrX* is not a conserved feature of specific clades, as for example *A. vitis* does not complement. Finally the functional diversification between FcrX and FliJ is also supported by the fact that, to the best of our investigations, FcrX has no role in the control of motility.

**Figure 5:**
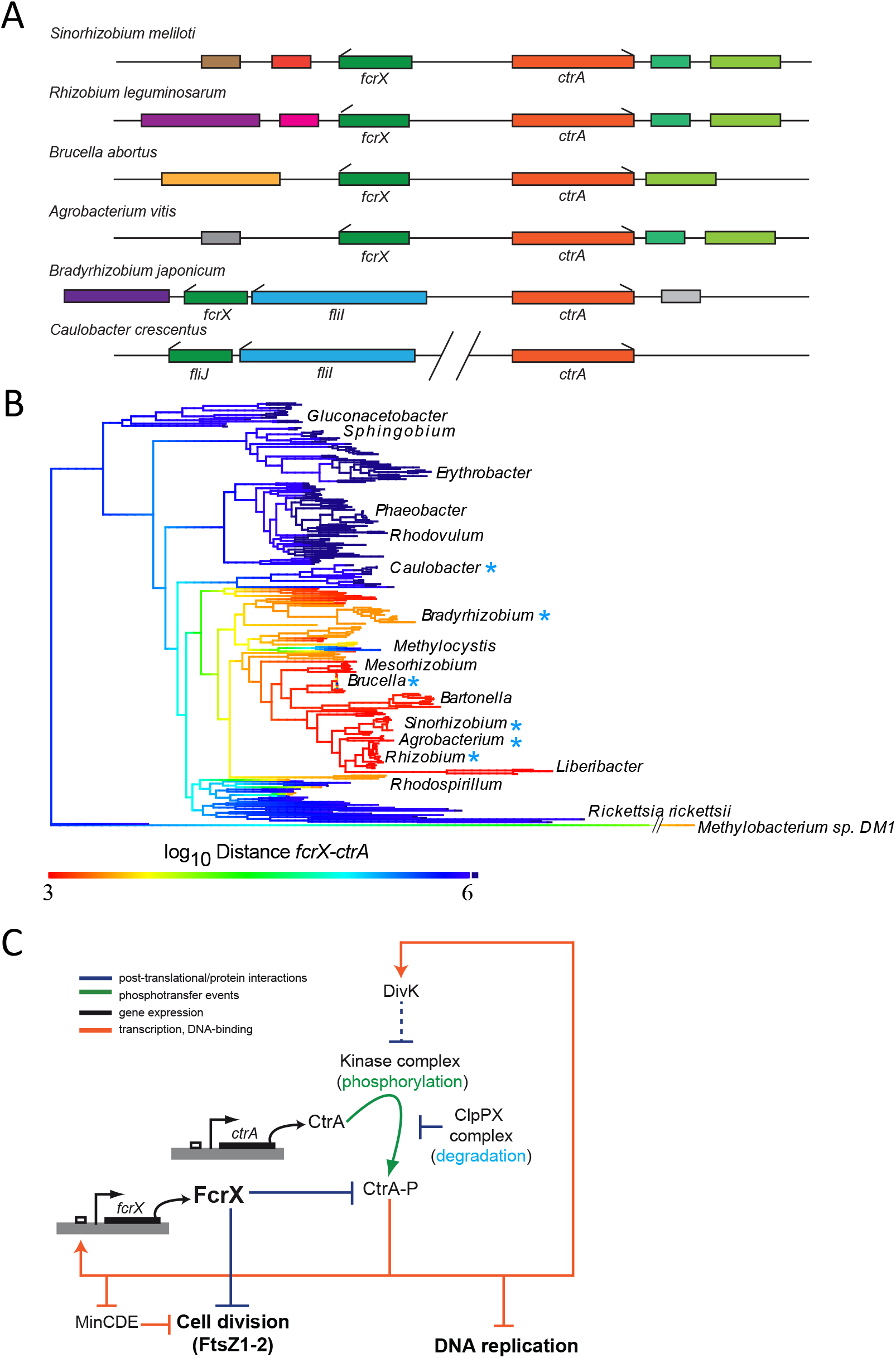
FcrX is conserved among alphaproteobacteria. (A) Typical organization of *fcrX* genomic loci in model alphaproteobacteria. The presence and the relationship between locations of *fcrX* and *ctrA* genes in those alphaproteobacterial species is highlighted with a blue asterisk in figure 5B. (B) Phylogenetic tree of FcrX orthologs-containing species (as described in Materials and Methods). The color code of the tree marks the distance between the *fcrX* and *ctrA* genes (values are bp). Species used in Figure 5A are marked with a blue asterisk, tree based on RecG sequences. (C) Model of FcrX role in cell cycle regulation with respect to main functions of cell cycle and CtrA/DivK/ClpXP essential regulators.

## DISCUSSION

In every organism, important functions of the cell are controlled by key factors (master regulators) coordinating many related elements. This is the case, for example, of the cell cycle in eukaryotes, based on cyclins ^32^, the sporulation process of *Bacillus subtilis*, controlled by Spo0A ^33^ and the cell cycle regulation in some alphaproteobacterial, such as *C. crescentus*,where it is controlled by the master regulator signal transduction protein CtrA ^34^. In this work we identified in *S. meliloti* a new one key factor of cell cycle and cell division that we named FcrX.

FcrX is a surprisingly small protein with no homology to other, so far characterized, regulators. We predicted that FcrX is an alpha helix-rich protein that can oligomerize, and shares ancestry with a previously characterized chaperon involved in flagellum physiology, named FliJ. In *C. crescentus*, this small protein is specifically required for flagellum functioning through the stabilization of another protein FliI, an ATPase involved in the export of flagellar subunits across the membrane using a dedicated type III protein secretion system ^35^. The analysis of synteny of FcrX orthologs clearly underlined this association between FcrX and FliJ-FliI, as many orthologs of FcrX are still organized in a *fliIJ* operon. Another striking feature of the analysis of the *fcrX* locus in many alphaproteobacteria is its proximity to the *ctrA* gene. As previously hypothesized, CtrA is considered as an ancestral flagellum regulator in many alphaproteobacterial species, including those in which CtrA also plays a role as cell cycle regulator ^4,36^. This conserved characteristic may suggest that FcrX evolved from FliJ, extending or changing its targets from flagellar components to the divisome protein FtsZ and the cell cycle regulator CtrA.

Indeed we showed here that FcrX binds directly CtrA in an unknown way and its presence plays a negative role on the steady state levels of CtrA. Conversely, CtrA in addition to many cell cycle genes ^10^ also controls *fcrX* transcription in the second half of DNA replication phase. From this point of view, CtrA-FcrX forms an essential negative feedback loop contributing to the oscillation of CtrA levels during cell cycle (Figure 5C). The presence of an essential genetic negative feedback loop has been also demonstrated in *C. crescentus*, in which DivK and CtrA are the main components of the feedback loop of this species ^8^. It is tempting to speculate that although the architecture of cell cycle regulation may change between different organisms, CtrA-DivK (linked by a transcriptional relationship) in *C. crescentus* and CtrA-FcrX in *S. meliloti*, the logical principles behind the regulation remain similar.

In addition, this CtrA-related crucial regulatory function of FcrX is not its only role, as this novel master regulator of cell cycle also negatively controls the main component of the divisome, FtsZ (FtsZ1 and FtsZ2 in *S. meliloti*), by direct protein-protein interaction. However, at this stage it cannot be excluded that FcrX interaction with FtsZ2 (both in BACTH and affinity columns) may involve a ternary complex with FtsZ1 (either from *E. coli* in BACTH or endogenous one in affinity columns) and FcrX, as FtsZ2 lacks a protein interaction domain found in FtsZ1. This dual activity of FcrX makes this factor a novelty in the knowledge of cell cycle regulation in alphaproteobacteria. Although cell division has been shown to be usually regulated by CtrA at the transcriptional level in *S. meliloti*, but also in *C. crescentus* or *B. abortus* ^34^, this is the first time that a negative regulator of cell cycle is able to connect directly to both cell cycle regulation and the divisome itself. This dual activity of FcrX is responsible of its severe depletion phenotype, leading to a block of cell cycle producing minicells that contain no DNA, but also keeping a mother cell with the genome able to produce new minicells. Considering potential applications that need minicells to perform specific functions ^37,38^, the depletion of *fcrX* represents a miniatured minicell factory, which may be exploited in the future for biotechnological purposes.

Another important aspect of FcrX functionality is related to symbiosis and bacteroid differentiation. It has been shown previously that bacteroids in *S. meliloti*, in order to become functionally mature, must eliminate CtrA and FtsZ, leading consequently to elongated cells with multiple copies of DNA. The discovery of a single regulator that is able to control negatively both CtrA and FtsZ suggests that FcrX may play a role during bacteroid differentiation. Indeed we have shown that FcrX is present in mature bacteroids and its function is required for a correct establishment of symbiosis. Accordingly, a strain constitutively expressing FcrX was able to increase plant biomass with respect to the wild-type situation, suggesting that promoting CtrA and FtsZ downregulation can increase the efficiency of the symbiosis, possibly by predisposing bacteria to terminal differentiation and making this process take place earlier rather than increasing it to a higher level. This will open new frontiers of sustainable agriculture by the use of improved bacterial inoculants based on FcrX deregulation. Even more interestingly, FcrX appears as a conserved factor in several rhizobial species further suggesting that this approach of plant growth amelioration may be extended to other agronomically-important legumes symbionts.

In conclusion, FcrX is a novel global factor controlling two essential key functions of the cell, regulation of cell cycle progression (CtrA) and cell division (FtsZ) (Figure 5C). This central position and its integrated role in a negative feedback loop with CtrA suggests that cell physiology may rely on FcrX regulation in order to perform higher levels of coordination of cell cycle. In the future, the investigation should move towards exploring how FcrX is regulated and what is the actual mechanism of CtrA and FtsZ1/2 inhibition by FcrX. FcrX indeed represents a small protein with capacities to interact with very diverse targets that may be a tool or a target for antibiotic therapies.

## MATERIALS AND METHODS

### Growth conditions

The strains used in this study are listed in the Table S3. *S. meliloti* 1021 and *E. coli* strains were grown in YEB medium (0.5% beef extract, 0.1% yeast extract, 0.5% peptone, 0.5% sucrose, 0.04% MgSO_4_·7H_2_O, pH 7.5) at 30°C and LB medium (1% tryptone, 1% NaCl, 0.5% yeast extract) at 37°C, respectively. Media were supplemented with appropriate antibiotics: Kanamycin (50μg/ml), Tetracycline (10μg/ml), Gentamicin (20μg/ml) for *E. coli*,Streptomycin (200μg/ml), Kanamycin (200μg/ml), Tetracycline (2μg/ml), Gentamicin (20μg/ml) for *S. meliloti*. Depletion strains of *S. meliloti* were grown in a medium supplemented with IPTG (100μM for *fcrX* and 1mM for *ctrA*); the depletion was accomplished by washing three times the culture and resuspending it in a medium lacking IPTG at OD_600nm_ = 0.3. The synchronization experiment was performed as described previously (De Nisco *et al*., 2014).

### Strain constructions

The two-step recombination procedure was used to perform the *fcrX* deletion using the integrative plasmid pNPTS138 as previously described (Pini *et al*., 2013). Deletions were verified by PCR using primers flanking the recombination locus (see primers in Table S4). To construct the fusion between the protein of interest and the fluorescent proteins (CFP or YFP), the Gateway procedure (Thermo Fisher) was used. First, the gene was amplified by PCR (see primers in Table S4) then introduced in the pENTR vector. Then, the vectors were mixed with the destination plasmid carrying the gene coding for the fluorescent protein, to perform the LR reaction as recommended by the manufacturer. The final product was amplified by PCR and cloned in the pSRK vector downstream the P*lac* promoter (Khan *et al*., 2005) and electroporated in *S. meliloti* as previously described ^39^.

To transduce the *fcrX::tetR* deletion the phage M12 was used ^40^. To do so, the bacteria were grown in LB containing 2.5 mM CaCl_2_ and 2.5 mM MgSO_4_ then mixed with the phage to give a multiplicity of infection of 0.5. The mixture was incubated at room temperature for 45 min and subsequently plated on LB plates with the appropriate antibiotics.

In order to identify the *fcrX* promoter, six different *PfcrX-fcrX* constructions were cloned into the *Plac* inducible plasmid pSRK and were electroporated in *S. meliloti* wild type cells. All *S. meliloti* clones containing an extra plasmid-encoded copy of *fcrX* with different promoter lengths were transduced using a phage M12 lysate produced from the strain Δ*fcrX::tetR* + *Plac-fcrX* and selected in presence or absence of IPTG. To verify the *fcrX* promoter sequence and its regulation by CtrA, the different constructions were introduced in pOTem1 ^26^ vector using the RF cloning procedure ^41^.

*S. meliloti* 1021 (wild type) Tn-seq data of the *fcrX* gene during growth in YEB medium was obtained from a previous study ^42^.

### Nodulation assays and analysis

*M. sativa* cultivar Gabès seeds were scarified with pure sulfuric acid for 8 min. After several washes with distilled water, the seed surface was sterilized with bleach (150ppm) for 30 min and seeds were washed again. Finally, seeds were soaked overnight under agitation in sterile water and then transferred onto a Kalys agar plate for one day at 30°C in the dark to allow the germination. The seedlings were planted in perlite/sand (2:1 vol/vol) in 1.5L pots in the greenhouse (24°C, photoperiod 16 h of light and 8 h of dark, humidity 60%) and were inoculated 7 days after planting with 50 ml per pot of the appropriate bacteria at OD_600nm_ = 0.05. Plants were watered every three days, alternating tap water and a commercial N-free fertilizer (Plant Prod solution [N-P-K, 0-15-40; Fertil] at 1 g per liter). Plants were harvested at 6 weeks post inoculation (42 dpi) to analyze bacteroid colonization and nodule development by confocal microscopy, level of bacteroid differentiation by flow cytometry and plant dry mass measurement ^43^.

### Electron microscopy

Bacteria were prefixed by adding an equal volume of fixative (2% glutaraldehyde in HEPES buffer 200mM, pH 7.2) to the culture medium. After 20 min, the medium was replaced by 1% glutaraldehyde in HEPES buffer for at least 1 h at 4°C. Bacteria were then washed with HEPES buffer, concentrated in 2% agarose (LMP Agarose, Sigma A9414), washed again with HEPES buffer and post-fixed in 1% osmium tetroxide (EMS 19150) for 1h at 4 °C. Samples were washed again in distilled water and treated with 1% uranyl acetate (EMS 22400) for 1 h at 4°C in the dark. Subsequently, samples were dehydrated in a graded series of acetone and embedded in Epon resin. Ultrathin sections (60–90 nm) were cut, stained with uranyl acetate and lead citrate and were analyzed using a Tecnai 200kV electron microscope (field electron interference or FEI). Digital acquisitions were made with a numeric camera (Oneview, Gatan).

### Confocal and wide field microscopy

Nodule imaging was performed on a SP8X confocal DMI 6000 CS inverted microscope (Leica) equipped with hybrid and PMT detectors, a 10x dry (Plan Apo + DIC (NA: 0.4, Leica)) and a 63x oil immersion (Plan Apo + DIC (NA: 1.4, Leica)) objectives. For each condition, multiple z-stacks were acquired (excitation: 405 nm; collection of fluorescence: 520-580 nm for calcofluor excitation: 488 nm; collection of fluorescence: 520-580 nm for Syto9 and excitation: 561 nm; collection of fluorescence: 520-580 nm for Propidium iodide). Stacks were transformed into maximum intensity projections using ImageJ software.

Time lapse experiments were performed on depleted cells deposited on agarose/YEB (with appropriate antibiotics and inducers) and observed every 10 minutes (up to 16h) on a Nikon Eclipse Ti E microscope equipped with a Yokogawa CSU-X1-A1 spinning disk system.

### qRT-PCR experiment

RNA was extracted from bacterial culture samples using Maxwell^®^ 16 LEV miRNA Tissue Kit (Promega). cDNA was produced using random hexamers as primers and the GoScriptTM Reverse Transcription kit from Promega. Amplification of 16S rRNA and *fcrX* cDNA was made using SsoFast EvaGreen Supermix 2X kit (Bio-Rad, France) on a CFX96 Real-Time System (Bio-Rad) instrument and the results were analyzed by Bio-Rad CFX Maestro version 1.1 software (Bio-Rad). For each sample, a biological duplicate was realized. Primers are listed in Table S4.

### Flow cytometry

Cells were heated 10 minutes at 70°C and then stained, depending on the experiment, with DAPI (300μM), Syto9 (2.5nM), Propidium iodide (2.5nM) and Potomac Gold (1mM). After 10 minutes of incubation at room temperature, the cells were processed with Cytoflex bench-top cytometer (Beckman-Coulter) and the data analyzed with CytExpert 2.5 software.

### Western blot experiment

The bacterial pellets were prepared and frozen at OD_600nm_ = 0.6. Western blot was performed as previously described (Pini et al., 2013). Anti-GroEL are commercial antibiodies against *E. coli* GroEL (Abcam). For anti-FcrX antibodies, *fcrX* was cloned into a pET derivative with N-terminal His6 tag using a Gateway cloning procedure as previously described (Skerker et al., 2005). His6-FcrX was purified on a nickel column and rabbits were injected using a 28 days protocol (Pini et al., 2013). Purified plasma was then used for western blots.

### Protein interaction experiments

For the Bacterial Two Hybrid experiment ^44^, the recommendations by the supplier (Euromedex) were applied. To construct recombinant proteins, vectors available from Euromedex were used. These vectors enable the in frame fusion of the proteins subunits of adenylate cyclase from *Bordetella pertussis* (T18, T25) at the C and N terminus. To test protein putative interactions, each appropriate combination of vectors was electroporated into the *E. coli* strain βHT101, deleted of the gene coding for the endogenous adenylate cyclase (*cya* strain). Positive control corresponds to the Tol-Pal from E. coli ^45^.

For biochemical protein-protein interaction analysis, a nickel affinity column was used. Cells of *E. coli* BL21 (D3A) expressing His6-FcrX and *E. coli* BL21 (D3A) with no expression vector were induced 3h with 100μM IPTG at 30°C. Cells were harvested, sonicated as previously described (Skerker et al, 2005) and soluble lysate of both strains was loaded onto prepacked nickel columns. After several washes of extraction buffer (Tris 100mM, NaCl 500mM, imidazole 30mM, pH 7.5), an *S. meliloti* sonicated lysate was loaded on the columns, washed as previously described, followed by elution at increasing concentrations of imidazole (5%, 10% and 30%), collecting the eluates. Samples were loaded on SDS-PAGE gels and analyzed by mass spectrometry (details in Table S1) or western blot using antibodies direct against CtrA and FtsZ1 or 2 (the same polyclonal antibody is able to detect both copies).

### FcrX conservation analysis

Homologs of RecG, FcrX and CtrA were identified by first blast ^46^ hits, with an e-value cutoff of 10^-4^. Proteomes for all alphaproteobacteria considered were downloaded from NCBI. Only complete genomes were considered for the analysis. Genomic distances were calculated by using coordinates in the corresponding “.gff” file; the distance dividing two genes was defined as the minimum distance in both directions, *i.e*. taking the circularity of the genome into account. Ancestral state reconstruction of distances was performed and mapped on trees with function contMap from R-package phytools ^47^. Alignments were performed with Muscle ^48^ and refined by hand in AliView ^49^; maximum likelihood phylogenetic reconstructions were performed with iqTree ^50^, with options -nt AUTO -alrt 1000 -bb 1000, which combines ModelFinder, tree search, ultrafast bootstrap and SH-aLRT test.

## ACKNOWLEDGMENTS

Sara Dendene and Quentin Nicoud were supported by a PhD fellowship from the Université Paris-Saclay. Shuanghong Xue benefited from a PhD fellowship from the Chinese Scholarship Council. The present work has benefited from the core facilities of Imagerie-Gif (http://www.i2bc.paris-saclay.fr), a member of IBiSA (http://www.ibisa.net), supported by France-BioImaging (ANR-10-INBS-04–01) and from the support of Saclay Plant Sciences-SPS (ANR-17-EUR-0007). We thank the IMM Transcriptomic facility for the RNA preparation and the qRT-PCR experiment; we also thank Artemis Kosta and Hugo le Guenno from the IMM Microscopy platform for Electron Microscopy acquisition and analysis. This work was funded by the Agence Nationale de la Recherche, grants no. ANR-17-CE20-0011 and ANR-21-CE20-0040. The authors thank Corine Foucault, Armelle Vigouroux, Solange Morera and Roza Mohammedi for technical help.

