## Supplemental figures for "*Sinorhizobium meliloti* FcrX coordinates cell cycle and division during free-living growth and symbiosis"

### FIGURE S1

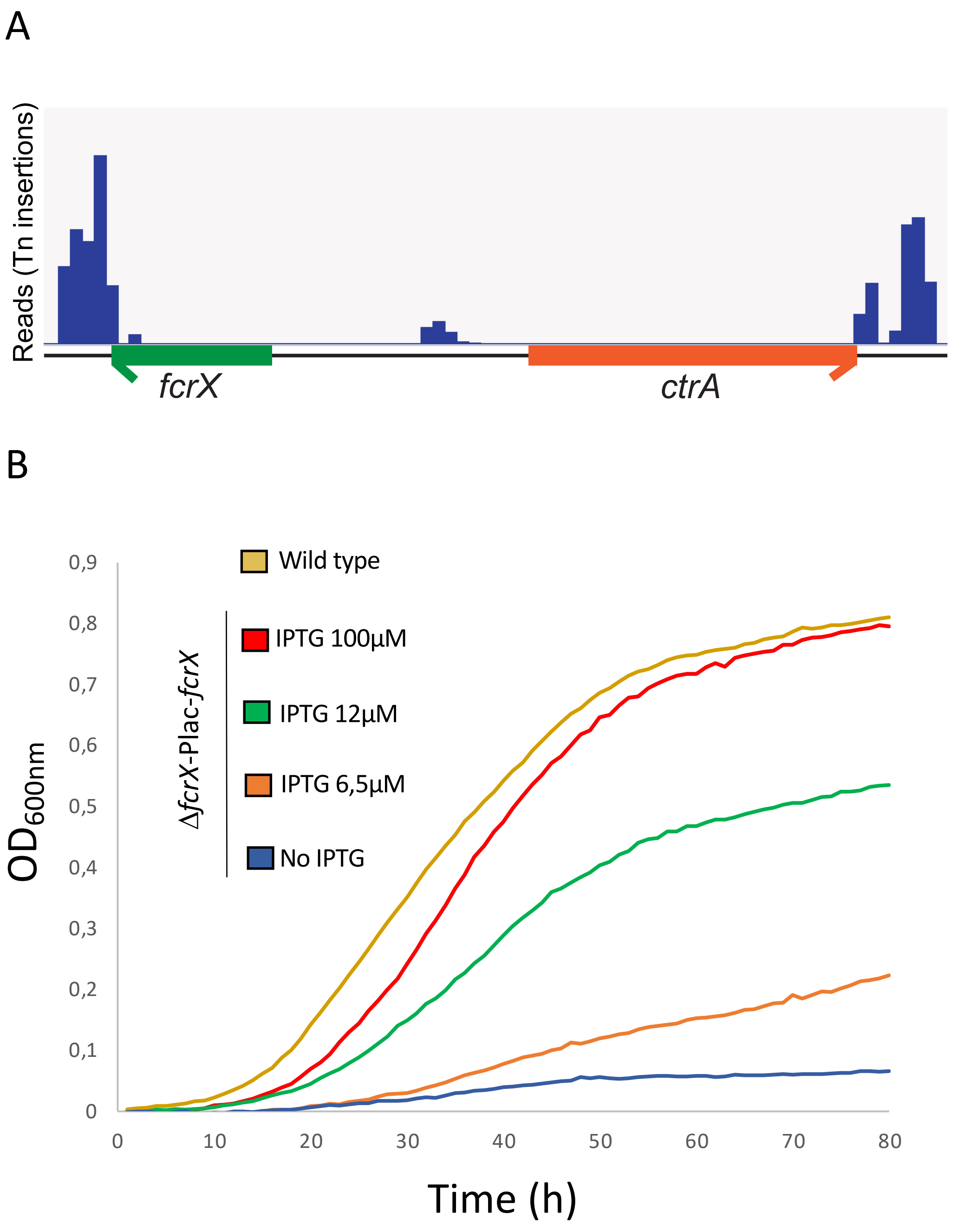


**Figure S1. Data supporting *fcrX* essentiality in *Sinorhizobium meliloti*.**

(A) Schematics of the *fcrX*-*ctrA* locus with Tn-seq insertion frequency.

(B) Growth curves of *S. meliloti* Δ*fcrX*-Plac-*fcrX* strain cultivated with different IPTG concentrations. Wild type strain was used as positive control.

### FIGURE S2

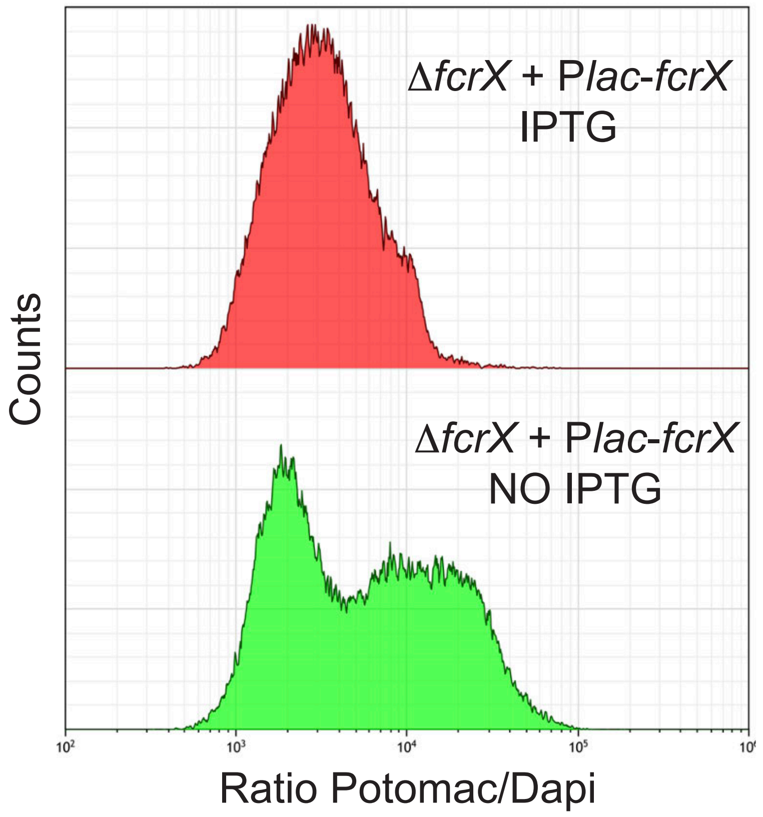


**Figure S2. Cells depleted of FcrX empty of DNA.**

Flow cytometry using Potomac (membrane) and DAPI (DNA) in Δ*fcrX*-Plac-*fcrX* with (red) and without IPTG (green).

### FIGURE S3

A

B

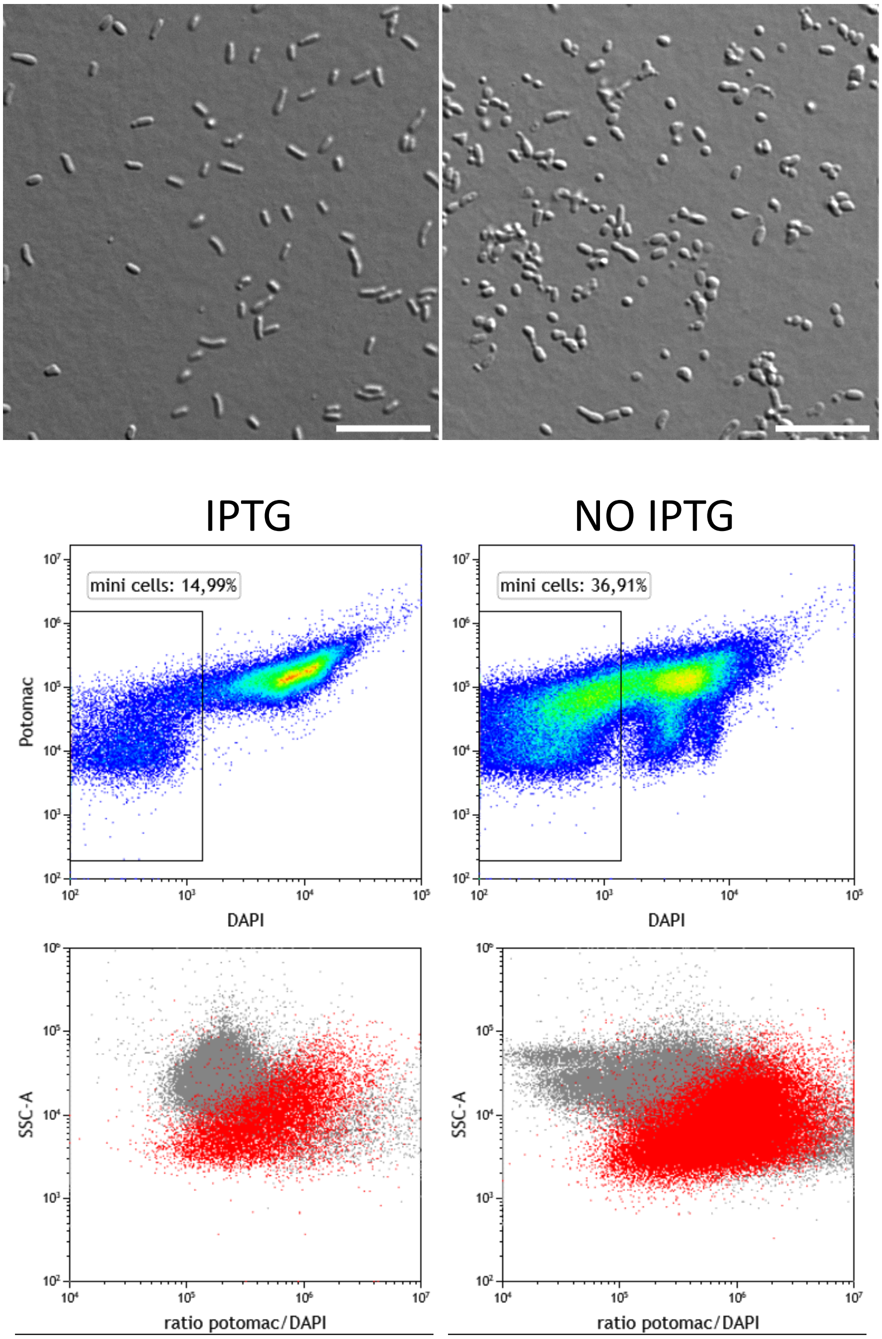


**Figure S3. Cells depleted of FcrX are minicells.**

(A) Transmission contrast microscopy of Δ*fcrX*-Plac-*fcrX* with (left) and without IPTG (right). Bars correspond to 10µm.

(B) Flow cytometry analysis using Potomac (membrane) and DAPI (DNA) in Δ*fcrX*-Plac-*fcrX* with (left) and without IPTG (right).

### FIGURE S4

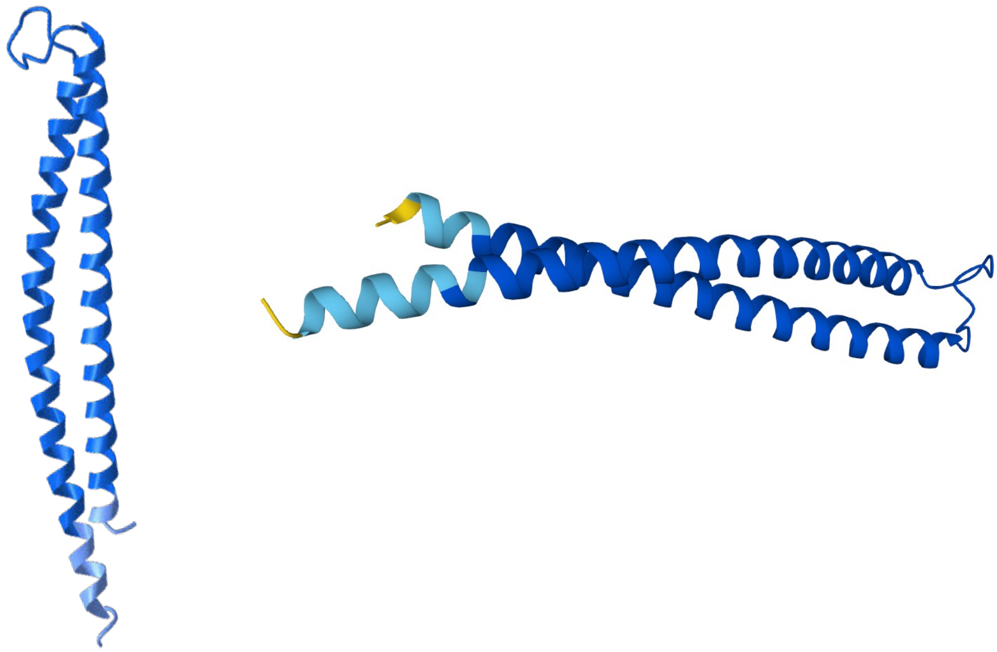

**Figure S4. Predicted structure of FcrX.**
Structure of FcrX (from two different angles) was predicted using Alphafold2. Darker blue corresponds to higher prediction confidence..

### FIGURE S5

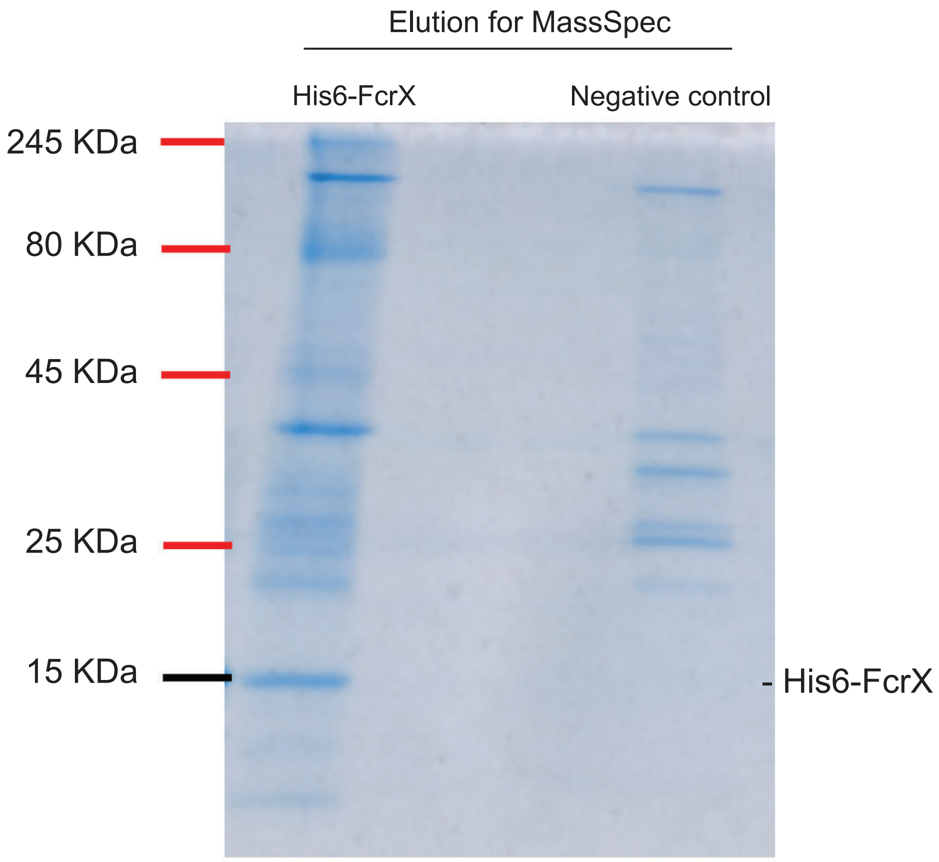


**Figure S5. Gel electrophoresis of HIS6-FcrX interacting proteins purified by affinity on a nickel column.**

Affinity column experiments were performed as described in Materials and Methods. His6-FcrX (left) and negative control (right) were eluted from the column using 30% imidazole buffer.

### FIGURE S6

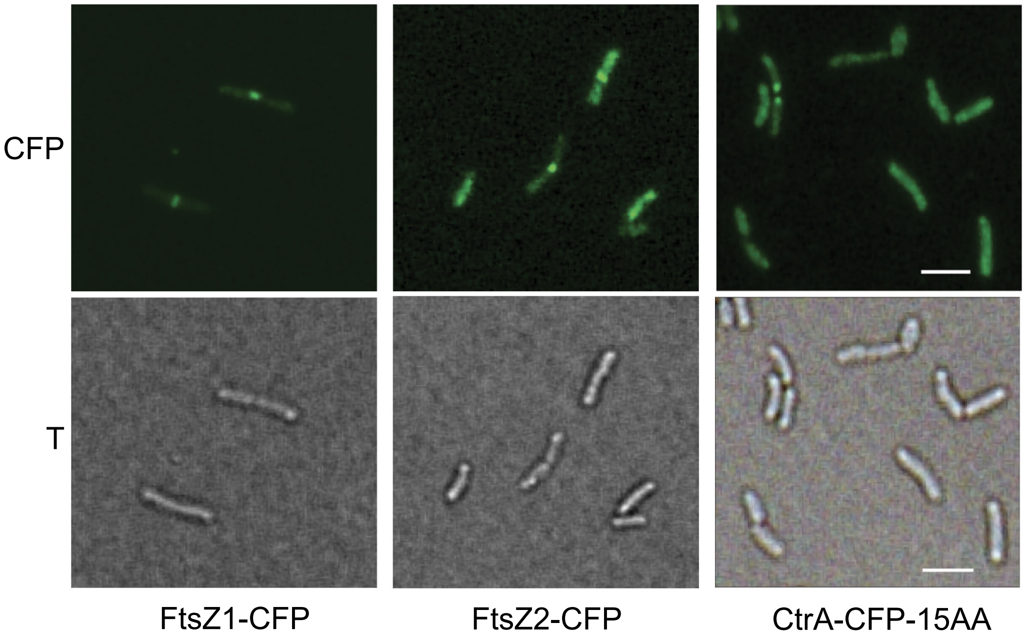


**Figure S6. Subcellular localization of FtsZ1, FtsZ2 and CtrA.**
C-terminal translational fusions of FtsZ1, FtsZ2 and CtrA with CFP were constructed as described in the text. Scale bar corresponds to 2 μm.

### FIGURE S7

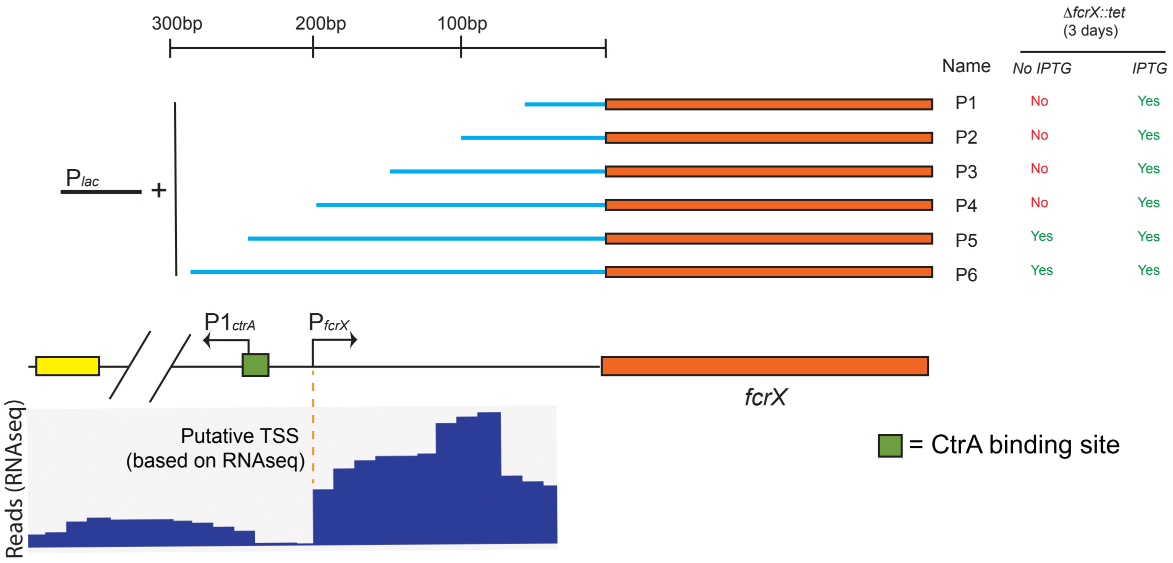


**Figure S7. Determination of *fcrX* promoter by deletion analysis.**
Several complementation regions were constructed with different length of the putative *fcrX* promoter , adding also a Plac promoter (inducible by IPTG). A transduction lysate containing the *fcrX* gene locus replaced by the tetracycline resistance cassette was transduced in the different *S. meliloti* backgrounds containing the different *fcrX* constructs (results of the transduction on the right table with and without IPTG). See text for more details. At the bottom, an RNA seq profile corresponding to the promoter region shows the coherence of RNAseq profile with the genetics presented here.

### FIGURE S8

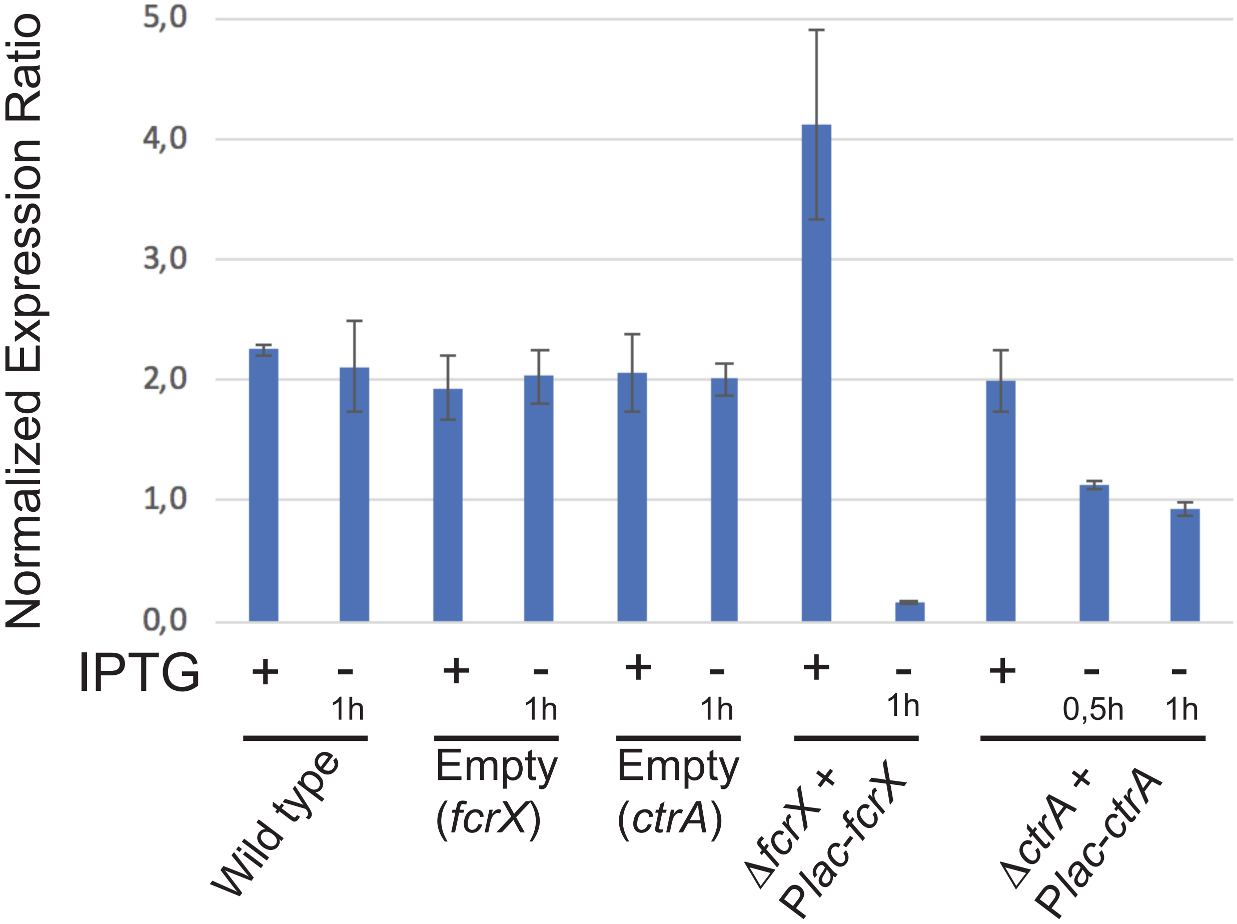


**Figure S8. Transcriptional levels *fcrX* in different genetic backgrounds.**
Expression transcriptional levels of *fcrX* normalized with respect to 16S were tested upon depletion of *fcrX* and *ctrA*. See text for more details.

### FIGURE S9

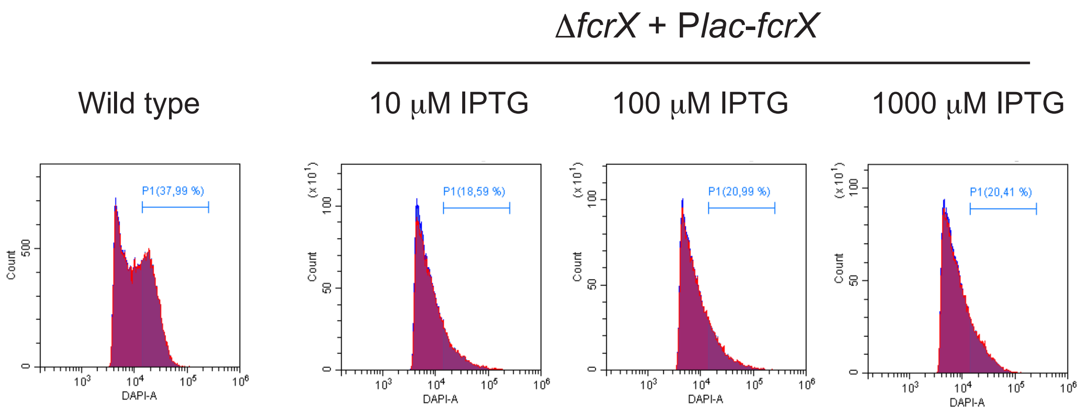


**Figure S9. DNA content of bacteroids extracted from 42 dpi nodules determined by flow cytometry as shown in figure 4A.**
Cells were labeled with DAPI.

### FIGURE S10

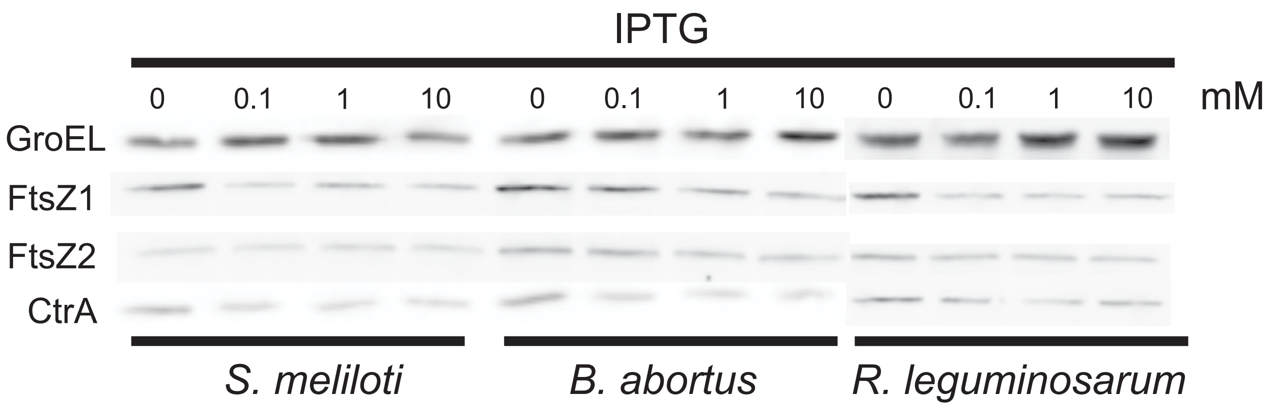


**Figure S10. CtrA and FtsZ1 and FtsZ2 steady state levels in *S. meliloti fcrX* depletion strains complemented by *S. meliloti, B. abortus* and *R. leguminosarum fcrXs.***
Western blots were performed as described in Materials and Methods.
